# Identification of domains in *Plasmodium falciparum* proteins of unknown function using DALI search on Alphafold predictions

**DOI:** 10.1101/2023.06.05.543710

**Authors:** Hannah Michaela Behrens, Tobias Spielmann

## Abstract

*Plasmodium falciparum*, the causative agent of malaria, poses a significant global health challenge, yet much of its biology remains elusive. A third of the genes in the *P. falciparum* genome lack annotations regarding their function, impeding our understanding of the parasite’s biology. In this study, we employed structure predictions and the DALI search algorithm to analyse proteins encoded by uncharacterized genes in the reference strain 3D7 of *P. falciparum*.

By comparing Alphafold predictions to experimentally determined protein structures in the Protein Data Bank, we found similarities to known domains in 353 proteins of unknown function, shedding light on their potential functions. The lowest-scoring 5% of similarities were additionally validated using the size-independent TM-align algorithm, confirming the detected similarities in 88% of the cases. Notably, in over 70 *P. falciparum* proteins the presence of domains resembling heptatricopeptide repeats, which are typically involvement in RNA binding and processing, was detected. This suggests this family, which is important in transcription in mitochondria and apicoplasts, is much larger in *Plasmodium* parasites than previously thought. The results of this domain search provide a resource to the malaria research community that is expected to inform and enable experimental studies.

## Introduction

*Plasmodium falciparum*, the malaria parasite responsible for over 600 000 deaths every year (World Health Organization, 2022), has a complex life cycle and its biology is still only partly understood. Of its 5268 predicted genes, 1407 remain without any annotation (release 62, January 2023 (Amos et al., 2022; Aurrecoechea et al., 2009)). Annotations of genes are most often based on sequence similarity to well characterized proteins in other organisms (Böhme et al., 2019). As part of an experimental gene-by-gene analysis of unknown proteins encoded on *P. falciparum* chromosome 3 we found that structural similarities of Alphafold structures matched well the experimental evidence of several proteins of unknown function (Kimmel et al., 2023). This indicates that many parasite proteins have a conserved evolutionary origin but evolved beyond recognition on the primary sequence level but not on the structural level. It can therefore be assumed that comparisons of the structural prediction of unknown proteins will yield useful information on proteins currently annotated as unknown. The availability of Alphafold structure predictions (Jumper et al., 2021) now enables the identification of domains based on geometric comparisons, independent of DNA and amino acid sequences. These similarities could provide information about gene products that so far lacked annotation. Here we apply this approach to all genes of unknown function in *P. falciparum*, specifically to the reference strain 3D7. Several sequence-independent algorithms exist to compare protein structures to each other based on geometry, including VAST (Gibrat et al., 1996), Dali (Holm, 2022), Foldseek (van Kempen et al., 2022), CE (Shindyalov & Bourne, 1998), the protein size-independent scoring algorithm TM-align (Zhang, 2005) and others (reviewed in (Kufareva & Abagyan, 2011)). For this screen we chose the DALI search algorithm as it has several advantages. It is set up to search the PDB and has an integrated graphical user (GUI) interface that also allows quick assessment of PFAM-annotated domains in the aligned structures. Further it has a built-in GUI for visual 3D-assessment of the alignment of hits to the query. We chose an open approach, where we screen all proteins of unknown function for the presence of any domains, rather than searching for a predefined set of domains or restricting the search to a small set of proteins of interest. This has the advantage that discoveries can be made independent of hypotheses that are related to specific processes in the parasite.

Using this approach, we here report the identification of domains with similarities to experimentally determined protein domains in 287 proteins encoded by genes of unknown function from *P. falciparum* 3D7 by an open-ended search. In addition, we used a targeted approach to search for proteins with similarity to the armadillo domain-contain ASA2 and ASA3 proteins which came to our attention as a frequent similarity in the open-ended approach. In this targeted search we found folds resembling heptatricopeptide repeats, which are typically involved in RNA-binding and -processing, in 53 previously un-annotated proteins, indicating that this family is much larger than previously though. We provide these results as a resource to the community to support experimental work.

## Results

### Identification of domains in proteins of unknown function by open-ended domain search

In order to identify known domains in *P. falciparum* proteins of unknown function, all 1407 proteins which were named protein of unknown function on PlasmoDB (The Plasmodium Genome Database Collaborative, 2001) were selected for analysis. The Alphafold predictions of the proteins encoded by these genes were compared to experimentally determined structures in the protein data bank (PDB) (Burley et al., 2019) using the search algorithm DALI (Holm, 2022). Included in the search were proteins that contained “globular” folds, thus excluding 476 structures which contained solely disordered, extended and fibrous folds, as this is a requirement for successful DALI searches. The resulting 931 hits were visually assessed to exclude structures that aligned only by two or fewer helices or beta strands and this resulted in the identification of similarity to at least one annotated domain in 287 of the 1407 unknown proteins (Figure 1A, Supplementary data 1).

**Figure 1:**
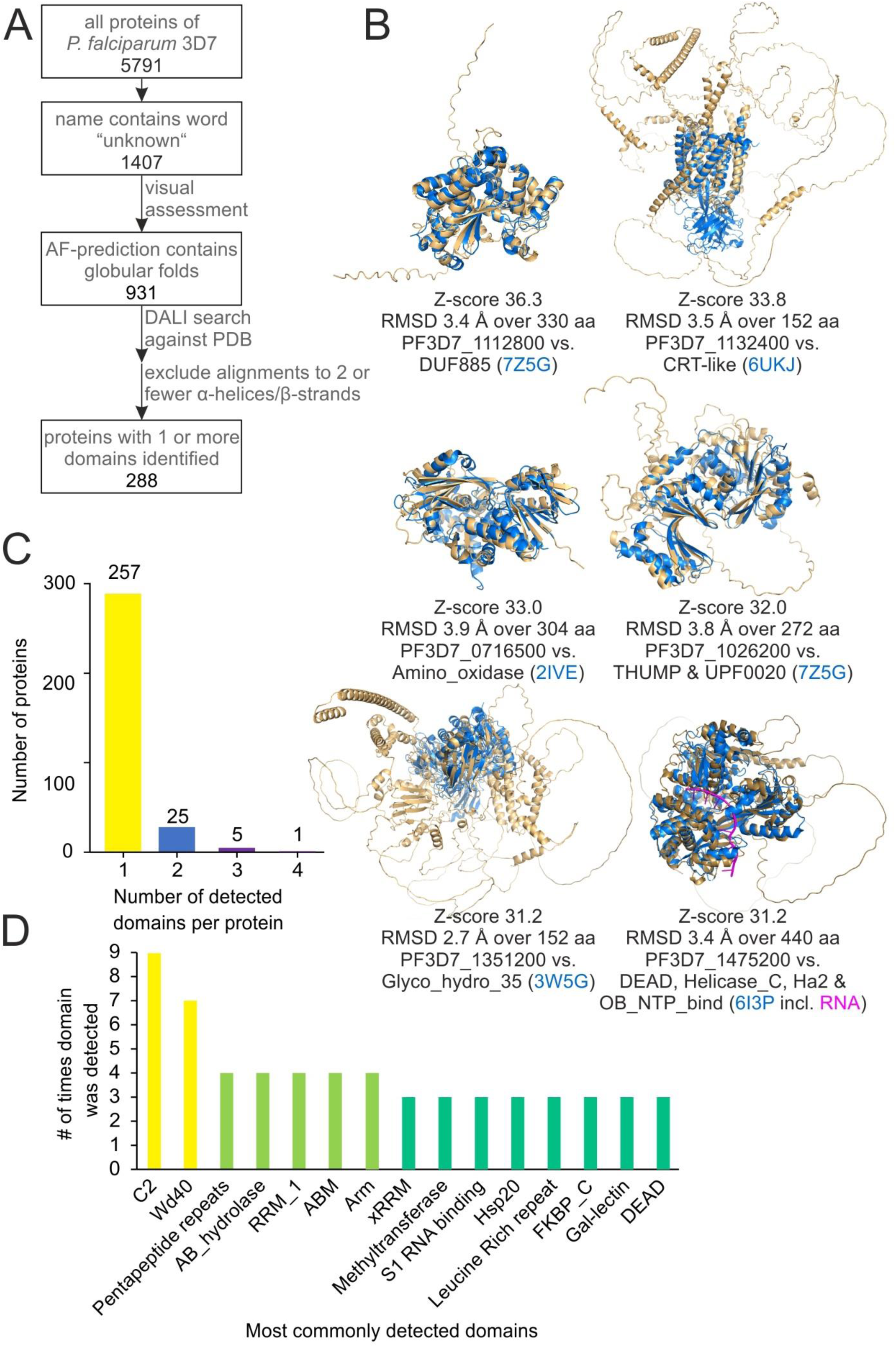
Identification of domains based on similarity of Alphafold prediction to PDB structures. (A) Pipeline for detection of domains in proteins of unknown function. (B) Alignment of Alphafold prediction (beige) to experimentally determined structured (blue) with domain annotation, with Z-scores >30. Indicated are the Z-score resulting from the DALI search, the root mean square deviation (RMSD) generated by Pymol cealign, the PlasmoDB gene accession number of the gene encoding the *P. falciparum* protein, the name(s) of the domain(s) in the experimentally determined structure according to Pfam and the PDB accession (blue) of the experimentally determined structure. (C) Number of proteins in which one, two, three or four domains were detected by the open-ended search. (D) Most commonly detected domains shown with number of times they were detected. Domain names as they appear in Pfam.

The DALI search assigns a Z-score to each alignment which reflects its quality, where Z-scores of below 2 indicate non-specific similarity. While all except one of the 287 structure alignments that were considered specific and a good fit by the visual inspection had Z-scores of 5 or higher, PF3D7_1336600 had a lower Z-score. However, the lower score in that case was due to its small size as it still showed a very good alignment (also shown in Figure 2). The six highest-scoring similarities had Z-scores of over 30 (Figure 1B). Of the proteins in which domains were identified, 254 contained one newly identified domain, 23 contained two and five proteins contained 3 newly identified domains (Figure 1C). As an example of a protein in which similarity to two domains was detected PF3D7_1013300 was visualized (Figure S1). Among the identified domains, most (>200) domains occurred only once, 27 occurred twice and 17 occurred three times or more (Figure 1D).

**Figure 2:**
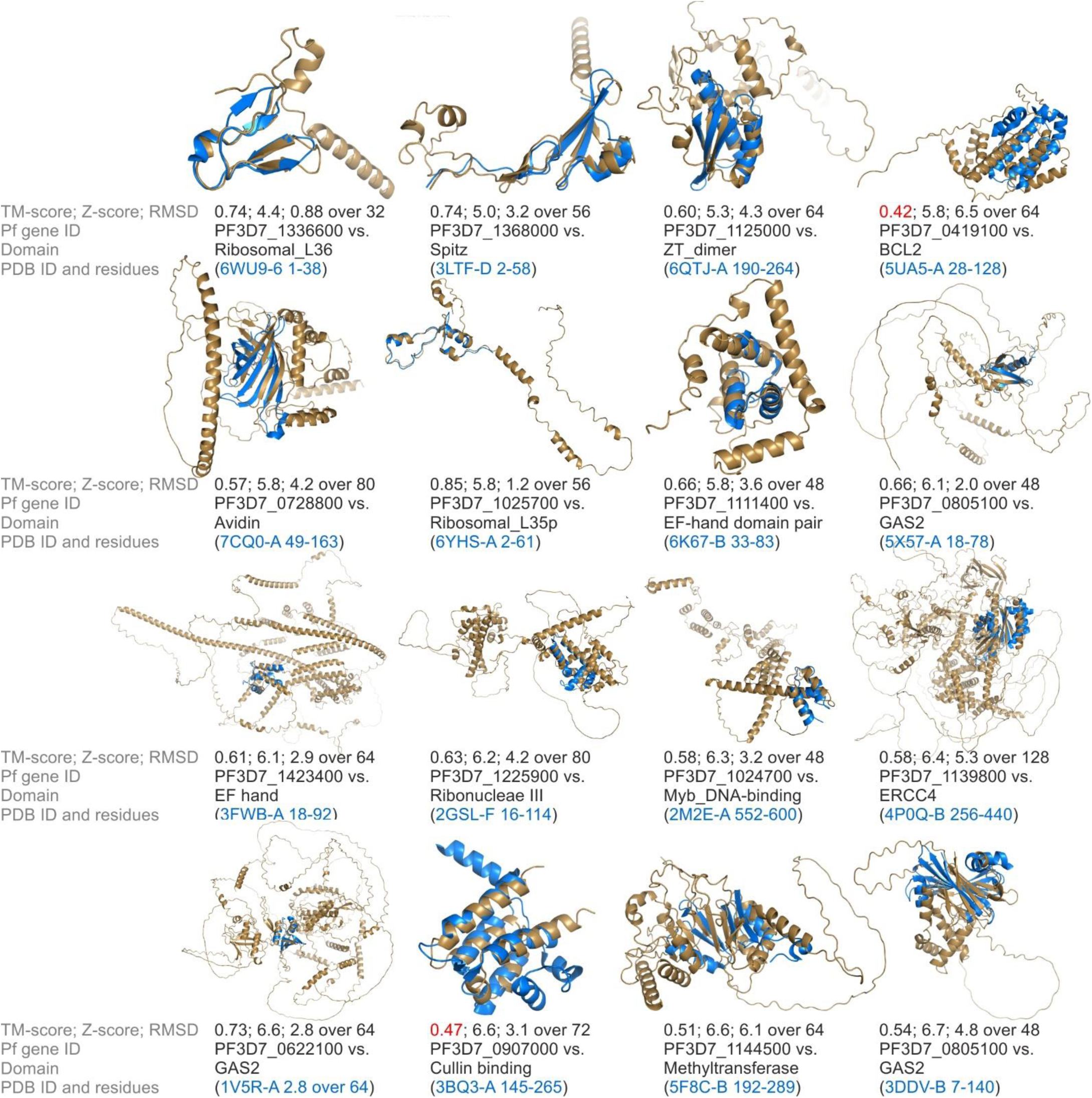
TM-align assessment of lowest-scoring 5% of aligned domains. The Alphafold structure of the *P. falciparum* protein (beige) is aligned to the indicated residues of the PDB structure (blue), which corresponds to the annotated domain. Given are TM-score generated by TM-align, Z-score generated by DALI, RMSD generated by Pymol cealign, PlasmoDB gene ID of *P. falciparum* gene that encodes the respective protein, name of the identified domain of similarity, the PDB ID of the structure that contained the domain and the amino acid residues at which the domain was found. For 3FWB-A residues 18 to 92 are shown even though the annotated domain only including residues 24-52 because structural similarity extended to this region. TM-scores under 0.5 are shown in red.

To validate these results, the lowest-scoring 5% of similarities found (based on DALI Z-score), were assessed using a second algorithm. For this TM-align was used which is independent of protein size (Zhang, 2005). This algorithm scores similarities between two protein structures on a scale from 0 to 1, where scores over 0.5 indicate that searched structures contain the queried fold. Of the 16 structures assessed 14 showed a TM-score over 0.5 (Figure 2), when compared to the top annotated hit from the DALI search (which was an experimentally determined structure downloaded from the PDB and cropped to the domain that showed similarity). The remaining two showed TM-scores over 0.4. Thus, the two algorithms show good agreement even among the lowest-scored similarities, suggesting a low proportion of false positive similarities.

### Comparison with published data

To assess whether the observed similarities have the potential to predict the function of the *P. falciparum* proteins of unknown function, we assess the literature for published experimental data. Although all of the 287 proteins for which similarity was found are encoded by genes that are annotated as of unknown function, some experimental data exists for eight of them (Table 1). The cellular location of the proteins encoded by PF3D7_0307600, PF3D7_0313400 and PF3D7_0319900 was previously assessed by fusing the endogenous gene with the sequence encoding GFP in an experimental screen of genes of unknown function from chromosome three in *P. falciparum* 3D7 parasites (Kimmel et al., 2023) (Table 1). PF3D7_0205600, PF3D7_1013300 and PF3D7_1329500 were analysed by the same approach (Schmidt et al., 2022) (Birnbaum et al., 2017; Khosh-Naucke et al., 2018) while the *P. berghei* ortholog of PF3D7_1132400 was expressed as an endogenous HA-fusion (Zuegge et al., 2001) and their cellular locations observed (Table 1). For PF3D7_0903600 functional data of the orthologue in *Toxoplasma gondii* recently became available (Dubois et al., 2023).

**Table 1:** Comparison of published data with predicted domains.

| PlasmoDB ID | Newly identified domain and its function | Experimental data available for protein | Match of domain identification and experimental data |
| --- | --- | --- | --- |
| PF3D7_0205600 | CDC45-like;<br>DNA replication<br>(Z-score 28.0) | GFP-fusion localizes to nucleus and cytoplasm (Birnbbaum et al., 2017). | Predicted function **supported** by experimental data. |
| PF3D7_0307600 | Rad51;<br>DNA repair<br>(Z-score 24.2) | GFP-fusion gives unknown localization pattern within the parasite (Kimmel et al., 2023). | Predicted function **not supported** by experimental data. Expected would be nuclear localization. |
| PF3D7_0313400 | AAA;<br>chaperone<br>(Z-score 13.5) | GFP-fusion localizes to the parasite cytoplasm (Kimmel et al., 2023). | Predicted function **possible** based on experimental data. |
| PF3D7_0319900 | UTRA;<br>Ligand-binding, modulates transcription factor activity in response to small molecules<br>(Z-score 6.7) | GFP-fusion gives unknown localization pattern in the parasite (Kimmel et al., 2023). | Predicted function **possible** based on experimental data. |
| PF3D7_0903600 | HOOK_N;<br>dynein-associated cargo adaptor proteins<br>(Z-score 12.0) | *Toxoplasma gondii* homolog TgHOOK regulates apical positioning and secretion of micronemes and contributes to egress, motility, host cell attachment, and invasion. Interacts with homologs of HOOK interaction partners TgFTS or TgHIP. (Dubois et al., 2023). | Predicted function **supported** by experimental data. |
| PF3D7_1013300 | Ncstrn_small (small lobe) and Nicastrin (large lobe);<br>Two domains making up nicastrin, a part of complex for intra-membrane proteolysis of integral membrane proteins. This protein is modified in Golgi or trans-Golgi network.<br>(Z-score 20.4) | GFP-fusion localizes to the periphery of ring stages, trophozoites and schizonts and is visible in ER in trophozoites and schizonts. (Khosh-Naucke et al., 2018).<br>Isolated from detergent-resistant membranes. HA-fusion gives “dotted labelling within the parasite cytoplasm” (Yam et al., 2013). | Predicted function **supported** by experimental data. |
| PF3D7_1132400 | CRT-like;<br>chloroquine resistance transporter and homologues. Arabidopsis homologues involved in thiol transport from plastid to cytosol.<br>(Z-score 33.8) | In *P. berghei* non-nuclear, non-apicoplast signal of HA-fusion protein. Predicted to have 9 to 10 TM domains. Western blot shows electrophoresis anomalies common to membrane proteins (Sayers et al., 2018). | Predicted function **possible** based on experimental data. |
| PF3D7_1329500 | TIG;<br>in cell surface receptors and in transcription factors for DNA binding<br>(Z-score 8.1) | GFP-fusion localizes to nucleus in ring and trophozoite stage, in addition foci in the parasite periphery at K13 compartment in trophozoites stage (Schmidt et al., 2022). | Predicted function **supported** by experimental data. |

PF3D7_0205600 and PF3D7_1329500 are located in the nucleus, supporting that the identified regions with similarity to CDC45-like and TIG domains, which are found in DNA replication proteins and transcription factors, respectively, might serve the same functions. PF3D7_1013300 showed similarity to both domains that make up nicastrins (Xie et al., 2014)and equally has a transmembrane domain and a short C-terminal tail (Figure S1). In other organisms nicastrin is part of the gamma secretase protein complex that proteolytically processes integral membrane proteins (Shah et al., 2005). Because inhibitors against the human gamma secretase complex had no effect on *P. falciparum* (Li et al., 2009) and homologs of the complex components were not found by sequence similarity, the gamma secretase complex was thought to be absent in *Plasmodium* parasites. Yet, the location of PF3D7_1013300 in the ER and the cell periphery would be consistent with nicastrin in other systems where it gets modified in the Golgi before transport to its final destination at the plasma membrane. Thus, the experimental data is compatible with the observed structural similarity and suggests that at least the nicastrin component of the gamma secretase complex is present in *P. falciparum* parasites.

For PF3D7_0903600 a HOOK_N domain-like fold was detected, which typically occurs in dynein-associated cargo adaptors. It’s ortholog in *Toxoplasma gondii* has recently been shown to interact with typical hook-interacting proteins and be involved in processes typical for hook proteins (Dubois et al., 2023), giving credibility to the detected domain.

For PF3D7_0313400 (AAA domain), PF3D7_0319900 (UTRA domain) and PF3D7_1132400 (CRT-like domain) the identified domains are not specific to distinct cellular locations but the experimentally observed cellular location do not disagree with their potential function. As such the domain predictions are possible based on the available experimental data. Solely for PF3D7_0307600, which we found to harbour similarity to a Rad51 domain (typically involved in DNA repair), the GFP-fusion protein was - contrary to our prediction - not found in the nucleus (Kimmel et al., 2023). This might be due to a faulty prediction, a repurposing in *P. falciparum* or an altered cellular location due to the GFP-fusion, which is possible given it is not essential (Kimmel et al., 2023; Schwach et al., 2015).

### Proteins containing Armadillo-like domains

It stood out that many unknown *P. falciparum* proteins showed similarity to Mitochondrial ATP synthase subunit ASA2 and Mitochondrial F1F0 ATP synthase associated 32 kDa protein ASA3. These two proteins are part of the Polytomella F-ATP synthase complex for which structures of several states were determined by single-particle cryo-electron microscopy (Murphy et al., 2019). As according to Interpro no domains are annotated in the ASA2 and ASA3 proteins, no equivalent domains in the structurally similar *P. falciparum* proteins could be annotated by our approach. To annotate this large group of *P. falciparum* proteins, we therefore sought to identify the shared domain of ASA2 and ASA3. Using their structures from PDB (PDB 6rd4), we performed a DALI search against all PDB structures. The top hits for either protein were ASA2 and ASA3, suggesting that both were very similar to each other. Further both were similar to proteins containing armadillo domains (PF00514), for example in catenin delta (PDB 3L6X (Ishiyama et al., 2010)), plakoglobin (PDB 3IFQ (Choi et al., 2009)), plakophilin (PDB 1XM9 (Choi & Weis, 2005)) and several other proteins, suggesting that both ASA2 and ASA3 contain domains, which are made up of several armadillo repeats.

We then performed DALI searches of these armadillo domains from ASA2 (residues 1-326) and ASA3 (full length) against all Alphafold-predicted structures of *P. falciparum* in the database. 1158 genes were found to encode proteins that contains domains similar to ASA2, ASA3 or both. As the Z-score is dependent on protein size, a suitable cut-off had to be determined for the common domain of ASA2 and ASA3. Spot checks of hits across different Z-scores were performed and showed that hits with a Z-score of 6.5 or higher showed good similarity to ASA2 or ASA3, with alignment of more than three armadillo repeats which each consist of one helix pair. To avoid false positives, an even more stringent Z-score cut-off of 7.0 was chosen which resulted in 121 *P. falciparum* (in 3D7) protein hits (Figure 3A, Supplementary data 2). All of them were found in both searches (ASA2 and ASA3). The hits included 18 heptatricopeptide repeat proteins, 13 RAP (**R**NA-binding domain abundant in **Api**complexans) proteins - both families are thought to be RNA-binding (Hillebrand et al., 2018; Hollin et al., 2021), - 3 that contain heptatricopeptide repeats and a RAP domain, 3 proteins that are named armadillo-domain containing proteins, 72 proteins of unknown function and 12 proteins with other annotations (Figure 3B). In 6 of these, there were also new domains identified in the open-ended DALI search shown in Figure 1, including DUF559, Importin-beta, Exportin 1-like, Armadillo domain and Atypical Armadillo domain. Both DUF559 domains (in PF3D7_0104600 and PF3D7_1207900) did not overlap with the identified armadillo domain, while the remaining four covered the same residues for which similarity to the ASA2-ASA3-armadillo domain was found (Figure S2).

**Figure 3:**
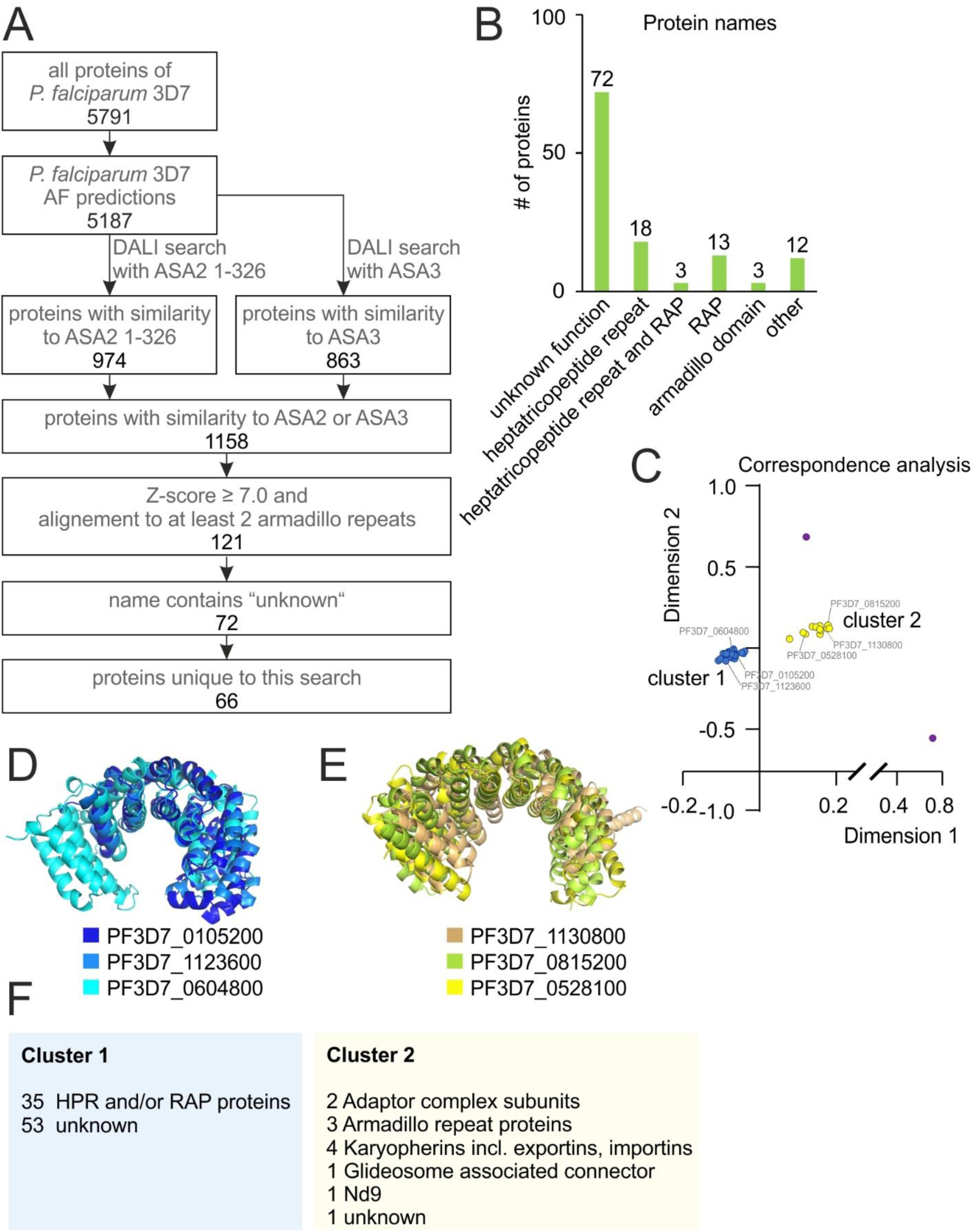
Identification of ASA2/3-like armadillo domains in *P. falciparum* proteins using Alphafold predictions. (A) Pipeline for detection of ASA2/3-like armadillo domains. (B) Annotations of *P. falciparum* proteins which were found to be similar to ASA2 and ASA3. (C) Correspondence analysis to cluster annotated *P. falciparum* proteins with similarity to ASA2 and ASA3 by structural similarity to each other. Dots corresponding to proteins shown in (D) and (E) are labelled. Purple dots do not belong to cluster 1 or 2. (D) Three representative structures selected from cluster 1 as defined in (C) aligned to each other using cealign in Pymol. (E) Three representative structures selected from cluster 2 as defined in (C). Structures shown in (D) and (E) were used to sort *P. falciparum* proteins of unknown function with similarity to ASA2 and ASA3 into clusters 1 and 2. (F) Number of proteins of different functions in cluster 1 and 2.

The amino acid sequences of these 121 proteins were aligned using Clustal Omega (Sievers & Higgins, 2018). The sequences of the proteins with annotation did not cluster according to their PlasmoDB annotations and generally showed little amino acid sequence similarity. We therefore decided to cluster them by structure. However, the 121 *P. falciparum* hits with similarity to ASA2 and ASA3 were too many to do this in one batch, thus first the Alphafold predictions of only those with an annotation in PlasmoDB were clustered using the DALI all-against-all algorithm. Two distinct clusters were observed and named cluster 1 and cluster 2 (Figure 3C). Cluster 1 was found to contain all heptatricopeptide repeat proteins, all RAP proteins and ARM2 (Figure 2C-F). All of these proteins are known or predicted to be RNA-binding proteins (Hillebrand et al., 2018; Hollin et al., 2021; Tang et al., 2019), while all proteins in cluster 2 are protein-binding proteins, some of which can also bind RNA (Frankel & Knoll, 2009; Fritz et al., 2009; Geiger et al., 2020; Henrici et al., 2020; Jacot et al., 2016; Jakel, 1998; Kumar et al., 2023; Xu et al., 2002). Three structures were randomly chosen to represent each cluster (Figure 3C-E) and aligned to batches of the 72 unknown *P. falciparum* proteins that harboured similarity to ASA2 and ASA3 (Figure S3). This assigned 53 new proteins to cluster 1 and one new protein to cluster 2, while 18 proteins could not be assigned to either cluster (Figure 3F).

Of the 20 proteins previously known to contain RAP domains, 5 were known to also contain heptatricopeptide repeats. Of the remaining 15 RAP proteins, 13 showed up in our search, suggesting that these also contain heptatricopeptide repeats. The DUF559 domains identified in two newly identified heptatricopeptide repeat-like proteins also show some similarity to RAP domains. Thus, the co-occurrence of RAP domains with heptatricopeptide repeats in the same protein seems to be a common combination.

## Discussion

In this study we used the DALI algorithm to search for similarities of the available Alphafold-predicted structures of all unknown *P. falciparum* (3D7) proteins to known domains in experimentally determined structures. This similarity search reduced the number of proteins without any annotation from 1407 to 1054, which is a 25% reduction. We expect the resulting set of several hundred identified domains to be a useful resource to the malaria research community that will help streamline further annotation and characterization efforts, and expected to aid the understanding of the molecular mechanisms of malaria parasites. It provides information on the potential function of these *P. falciparum* protein inferred from the structural similarity to known proteins in other organisms. Additionally, in combination with functional and interactome data these domain searches can aid and substantiate the assignment of proteins to protein groups or complexes which together serve a specific function as e.g. recently done for proteins in mitosis (Kimmel et al., 2023, Brusini et al., 2022).

Open-ended approaches have the advantage that they can discover unexpected components and pathways. The search in this study gives hints to the unexpected presence of nicastrin, which is part of a functionality thought to be absent in *Plasmodium* parasites. This approach can also provide hints for proteins that were expected to be present but had so far not been identified. One example from our dataset is PF3D7_0404300 which shows similarity to Ran-binding proteins and could be a missing component in transport in and out of the nucleus.

Unexpectedly, we discovered a large group of proteins in the parasite with very similar domains, including all known heptatricopeptide repeat and RAP proteins as well as 53 unknown proteins. Proteins containing heptatricopeptide repeats are known to bind and process polycistronic precursor RNAs into mRNAs and rRNAs for the mitochondrial ribosomes and to stabilize mRNAs to prevent decay (Hillebrand et al., 2018). Heptatricopeptide repeats proteins contain 37 amino acid long repeats, while proteins from the related families of pentatricopepide repeat proteins and octatricopeptide repeats proteins contain repeats which are 35 and 38 amino acids in length. Hepta-, penta- and octatricopeptide repeat proteins perform comparable RNA binding and processing functions, yet their distribution among different organism groups, like plants, apicomplexans and animals, varies (Hillebrand et al., 2018). In humans only 6 heptatricopeptide repeat proteins have been detected while around 70 were predicted in plants and dinoflagellates. In plants this is in addition to 450 pentatricopepide repeat proteins. In addition to the 18 previously described heptatricopeptide repeat proteins in *Plasmodium*, we here identified many more potential members of this group. If all 54 unknown proteins in cluster 1 of our clustering of the ASA2 and ASA3 hits indeed are such RNA-binding proteins, this would place the number of heptatricopeptide repeat-like RNA-binding proteins in *Plasmodium* (72) in a similar range as in plants. Interestingly, while some the *P. falciparum* proteins with similarity to heptatricopeptide repeat proteins contain insertions that break the 37-amino acid repeat motif that gives this domain its name, the geometry of the 37-amino acid repeat seems conserved.

Another interesting aspect is that we found heptatricopeptide repeat-like folds in a large proportion of proteins containing RAP and RAP-like DUF559 domains. RAP domains consist of a restriction endonuclease-like fold and are found in RNA-binding proteins (Hollin et al., 2021). Thus, heptatricopeptide repeats and RAP domains seem to be a common combination, suggesting that the two domains might act in concert.

While we believe this search to provide a valuable and useful resource, there are a number of limitations. The first inherent limitation is that this study used predicted structures of *P. falciparum* proteins rather than experimentally determined structures, which should not be viewed with the same confidence as experimentally determined structures. This circumstance has been discussed extensively following the introduction of Alphafold (Jones & Thornton, 2022; Subramaniam & Kleywegt, 2022; Terwilliger et al., 2022).

A second limitation was that the search presented here was not exhaustive. To the predicted structures, we applied DALI, the only search platform with which it was feasible to manually analyse several hundred protein structures in a reasonable time. This throughput is largely possible because of the built-in alignment and domain annotation viewers. The DALI algorithm scores search results based on length and quality of the structure alignment of the whole query structure.

As a result, this algorithm favoured the discovery of one domain of highest confidence, while further domains in the same protein were only discovered if they incidentally had similar scores as the first domain or if they occurred in the same experimentally determined structure that the query aligned to. This is one reason, why the search presented in this study was not exhaustive. Other search algorithms could circumvent this drawback, for example the VAST algorithm first detects potential domains in the query structure and then performs searches for each of them individually in addition to the search with the complete query structure. Generally, the use of other search algorithms, such as VAST (Gibrat et al., 1996), Foldseek (van Kempen et al., 2022) and PDBeFold (Krissinel & Henrick, 2004), might result in the discovery of different domain similarities due to their different implementations and ways of scoring similarity. This search being not exhaustive is an important limitation that should be kept in mind when using these findings. Nevertheless, some of the identified domains already give clear functional indications and a protein of interest can then be analysed in more detail using other algorithms in which case further domains might be identified.

A third limitation of our approach is that domains that were not annotated in experimentally determined structures could not be found. Searching against predicted structures of well-studied and annotated proteins might therefore expand the number of detected domains. Of course, with the caveat that predicted structures harbour some uncertainty.

Finally, the identified domains were not all computationally and none were experimentally validated in this study. Computational validation could be achieved by re-assessing the detected similarities using a second algorithm. This could for instance be the cealign command in Pymol (Figure 1B) followed by confirming or rejecting annotations based on a cut-off score, as we have done previously (Schmidt et al., 2022). It could also be done with the TM-align algorithm which has a pre-defined cut-off (Zhang, 2005) and was here used to validate the lowest 5% of the hits from this work. This confirmed 14 of 16 hits, overall giving a good credibility for the detected similarities, particularly as it can be assumed that the hits with higher scores are more reliable. Experimental validation is the gold standard for any structure prediction (Sanderson & Rayner, 2017). The assessment of available data in published literature on eight proteins annotated here, suggests that the majority of similarities found might align with the function that can be inferred from the domain similarities. However, a case-by-case validation is the only way to know whether the domains were identified correctly and whether these domains serve the same function in *P. falciparum* as in other organisms. We did not embark on a systematic experimental validation as this would take a considerable amount of time and we believe that rapidly providing these search results to the wider community will be more beneficial.

In conclusion, the results presented here are an example for the power of sequence-independent structure comparison approaches for reducing the number of genes lacking annotation and information on their potential biological function. Here applied to *P. falciparum*, it sheds light on the biology of the malaria parasite and provides starting points for future research which will lead to a better understanding of the biology of this pathogen.

## Methods

### Open-ended search

Lists of genes were downloaded from PlasmoDB v61 based on genomic location. All genes which contained the word “unknown” in the gene name, for example “conserved Plasmodium protein, unknown function”, were included in the analysis. Pseudogenes were excluded. Available Alphafold (alphafold.ebi.ac.uk, accessed December 2022-Mai 2023) (Jumper et al., 2021) structures were visually assessed for a globular and compact fold which is a prerequisite for a successful DALI search (Holm, 2022). The suitable Alphafold structures were submitted to a DALI PDB search as available in the web application (http://ekhidna2.biocenter.helsinki.fi/dali/, accessed December 2022-Mai 2023). The matches against PDB90 were analysed. The hits with the highest Z-scores were visually assessed for suitable alignment (excluded alignment of just 2 or fewer α-helices or β-strands) in the DALI 3D visualization tool. Of the top seven to ten hits with good alignment, the Pfam 35.0 (Mistry et al., 2021) domain annotations were viewed in the DALI PFAM tool. A domain annotation for a region of a PDB hit that aligned to the query Alphafold structure was considered a reliable hit if there were no conflicting domain annotation. Domain annotations were considered conflicting if other proteins with differently annotated domains aligned to the same amino acid residues and had a Z-score within 1.0 of the protein harbouring the domain of interest.

### Validation with TM-align

The 5% of domains with the lowest Z-score resulting from the open-ended DALI search were further validated. For each of these, the PDB structure of the top hit of the DALI search was cropped to only contain the residues annotated as the domain for which similarity was found. Where the top hit did not contain this annotation, the highest-scoring PDB structure with this annotation was used. The Alphafold structure was compared with this cropped PDB structure using TM-align (Zhang, 2005) as available in the web application (https://seq2fun.dcmb.med.umich.edu//TM-align/, accessed December 2022-Mai 2023). The TM-score based on the length of the cropped PDB structure was assessed.

### ASA2/ASA3 search

All protein structures predicted for *P. falciparum* by AlphaFold (Jumper et al., 2021) were searched using ASA2 residues 1-326 and full length ASA3 (chain 2 and 3 of PDB 6rd4 (Murphy et al., 2019)) using the DALI AF search (Holm, 2022). Alignments were visually assessed for at least 3 aligned armadillo repeats (helix pairs), resulting in all proteins with a Z-score larger or equal to 7.0 to be included for further analysis.

### Visualization and further assessment

Protein structures were analysed and visualized using PyMol 2.4.0 (Schrödinger, USA). Alignments were performed using the PyMol command cealign. Figures were arranged in CorelDraw X6-8.

## Acknowledgements

We thank Jesper Del Messier, Ana Ivan, Jan Stephan Wichers-Misterek and Isabelle Henshall for helpful discussions. This research made use of PlasmoDB, a VEuPathDB resource (Amos et al., 2022). HMB acknowledges funding by the Ortrud Mührer Fellowship of the Vereinigung der Freunde des Tropeninstituts Hamburg e.V.. HMB and TS acknowledge funding by the European Research Council (ERC, grant 101021493).

---

**Supplementary Figure 1:**
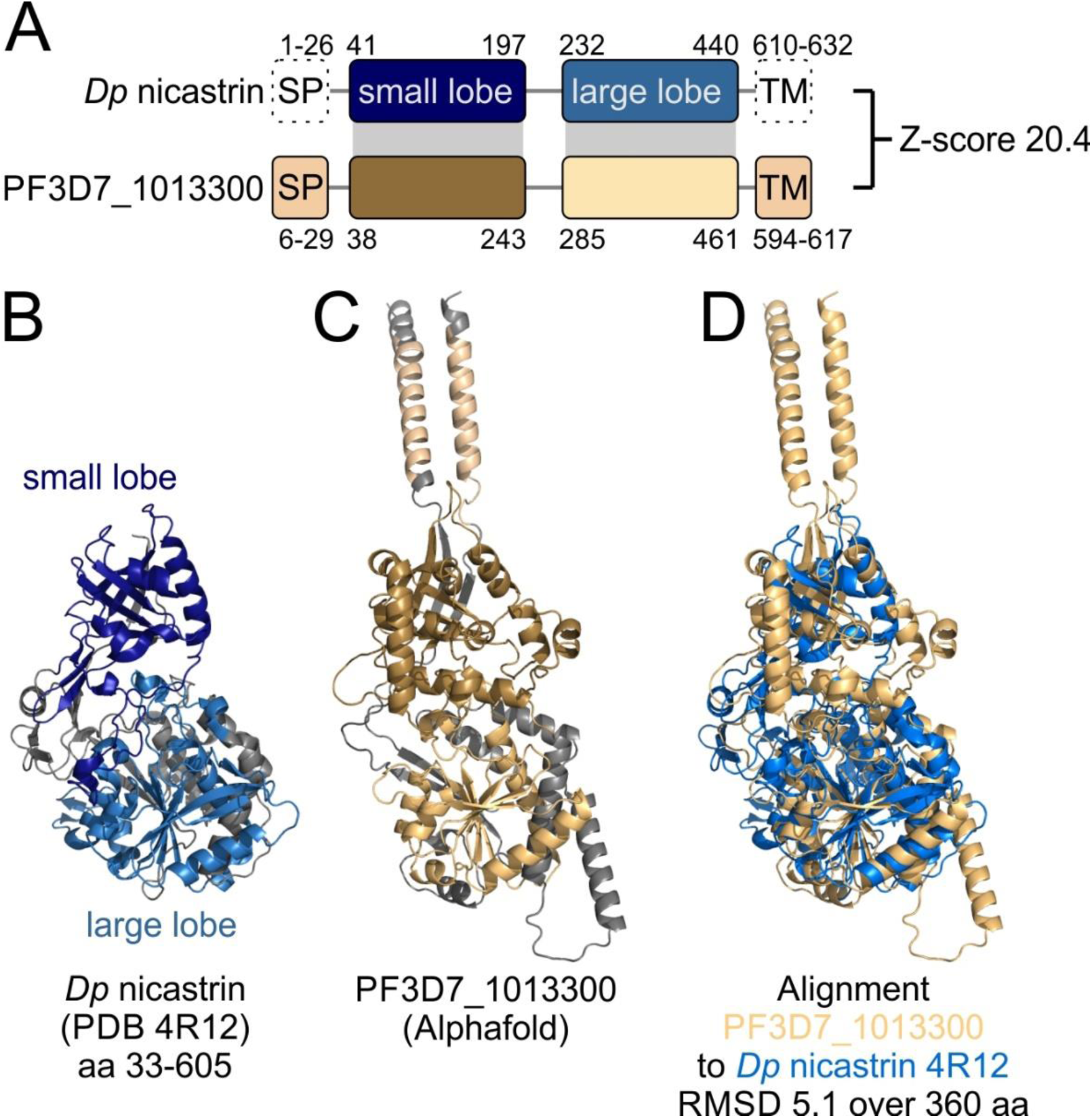
Nicastrin domains detected in PF3D7_1013300. (A) Schematic representation of *Dictyostelium purpureum* nicastrin and PF3D7_1013300 (not to scale). Grey elements show corresponding domains. Z-score resulting from DALI score is indicated. Secondary structure elements of *D. purpureum* nicastrin that are not present in the crystal structure PDB 4R12 are shown with dashed lines. SP, predicted signal peptide; TM, transmembrane domain. (B) Crystal structure of *D. purpureum* nicastrin (PDB 4R12). Residues coloured as in (A). (C) Alphfold structure prediction of PF3D7_1013300. Residues coloured as in (A). (D) *D. purpureum* nicastrin (PDB 4R12, blue) and Alphfold structure prediction of PF3D7_1013300 (beige) aligned to each other using cealign in Pymol. RMSD generated by Pymol cealign is indicated.

**Supplementary Figure 2:**
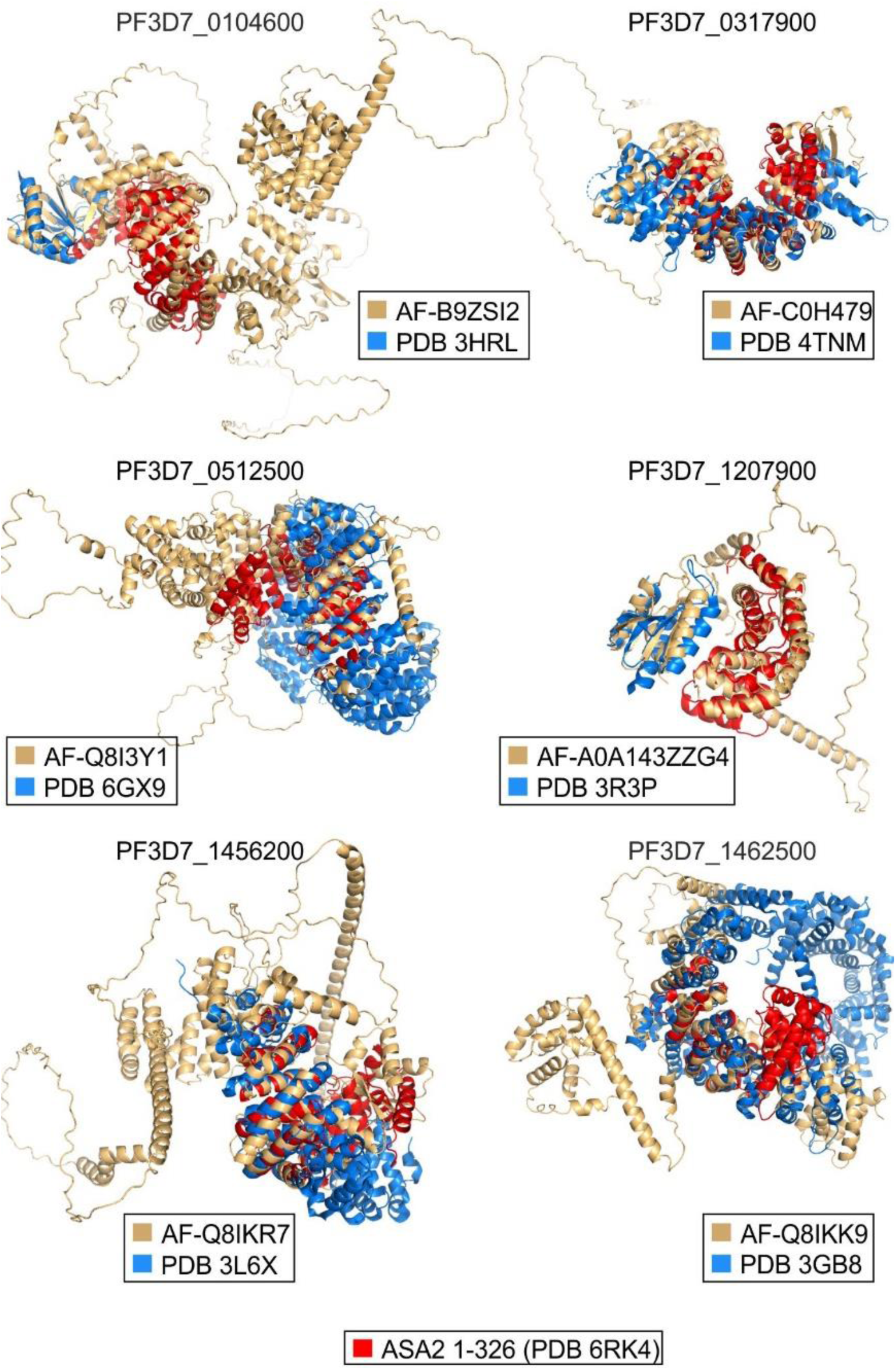
Spatial overlap of ASA2 with hits identified in open-ended domain search. Alphafold structure predictions of proteins for which domains were found in the open-ended search and in the ASA2/ASA3-based search are shown (beige), and aligned to ASA2 residues 1-326 from PDB 6RK4 (red) and to the highest-scoring domain-annotated hit of the open-ended search (blue). For PF3D7_1207900 only ASA2 residues 1-170 are shown for clarity. For PF3D7_0104600 and PF3D7_1207900 the domains from the open-ended search (both DUF559, blue) and the armadillo domain (red) align to different regions of the target protein. For the other four structures the domains align to the same region of the target protein.

**Supplementary Figure 3:**
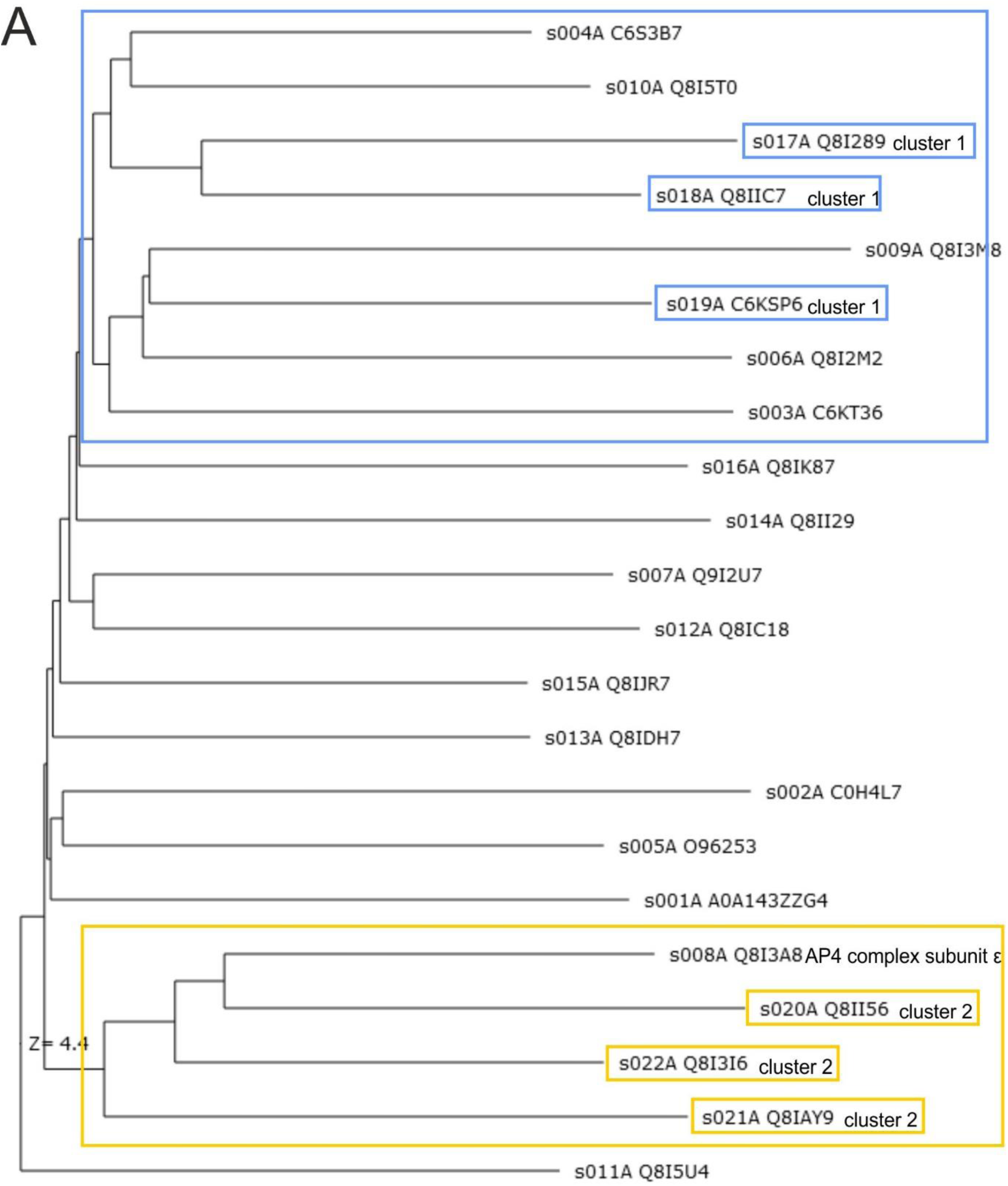
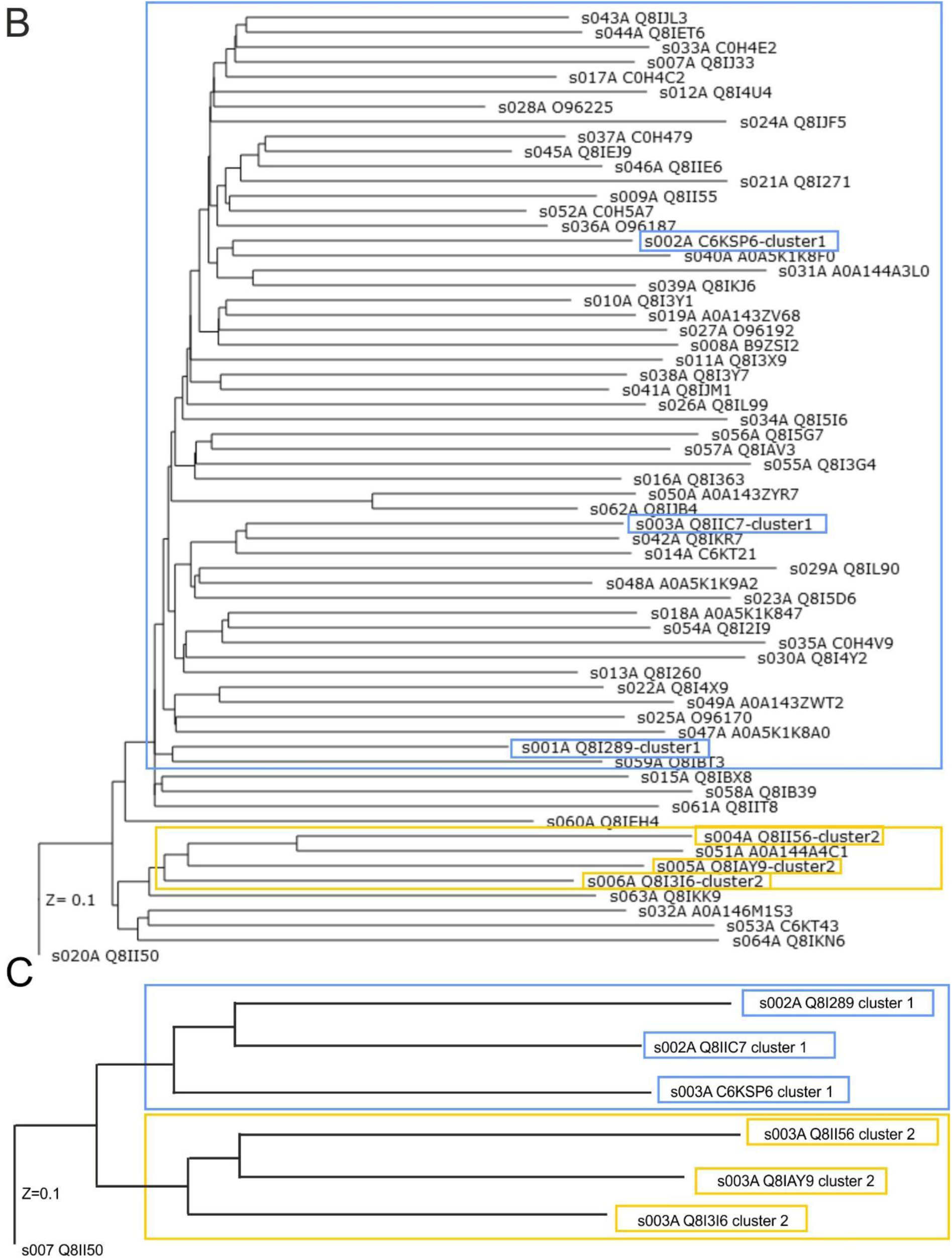
Structure based clustering of unknown proteins with ASA2/ASA3-like armadillo domains by DALI all-against-all search. Proteins of known functions which are representative of cluster 1 are labelled and highlighted in blue, those representative of cluster 2 are labelled and highlighted in yellow. Proteins considered to belong to the same cluster as the representative proteins are framed in the same color. (A) Batch 1. (B) Batch 2. (C) Batch 3.

